# Pangenomics to understand prophage dynamics in the *Pectobacterium* genus and the radiating lineages of *P. Brasiliense*

**DOI:** 10.1101/2024.09.02.610764

**Authors:** Lakhansing A. Pardeshi, Inge van Duivenbode, Michiel J. C. Pel, Eef M. Jonkheer, Anne Kupczok, Dick de Ridder, Sandra Smit, Theo A. J. van der Lee

## Abstract

Bacterial pathogens of the genus *Pectobacterium* are responsible for soft rot and blackleg disease in a wide range of crops and have a global impact on food production. The emergence of new lineages and their competitive succession is frequently observed in *Pectobacterium* species, in particular in *P. brasiliense*. With a focus on one such recently emerged *P. brasiliense* lineage in the Netherlands that causes blackleg in potatoes, we studied genome evolution in this genus using a reference-free graph-based pangenome approach. We clustered 1,977,865 proteins from 454 *Pectobacterium* spp. genomes into 30,156 homology groups. The *Pectobacterium* genus pangenome is open and its growth is mainly contributed by the accessory genome. Bacteriophage genes were enriched in the accessory genome and contributed 16% of the pangenome. Blackleg-causing *P. brasiliense* isolates had increased genome size with high levels of prophage integration. To study the diversity and dynamics of these prophages across the pangenome, we developed an approach to trace prophages across genomes using pangenome homology group signatures. We identified lineage-specific as well as generalist bacteriophages infecting *Pectobacterium* species. Our results capture the ongoing dynamics of mobile genetic elements, even in the clonal lineages. The observed lineage-specific prophage dynamics provide mechanistic insights into *Pectobacterium* pangenome growth and contribution to the radiating lineages of *P. brasiliense*.

## Introduction

In plants, soft-rot disease is mainly caused by bacteria from the *Pectobacterium* and *Dickeya* species, belonging to the *Pectobacteriaceae* family. *Pectobacterium* spp. is among the top 10 plant pathogens and infects potatoes, ornamental plants and vegetable crops (*1*). Although a number of *Pectobacterium* species, such as *P. brasiliense*, *P. atrosepticum*, *P. versatile*, *P. parmentieri*, *P. polaris*, and *P. punjabense* cause soft-rot disease in plants (*2*), *P. brasiliense* is one of the dominant species, known to infect at least 19 different plant species across the world (*3*). The *P. brasiliense* was first detected and characterized as a pathogen associated with blackleg (BL) disease of potato (*Solanum tuberosum L.*) in Brazil and initially named *Erwinia carotovora subsp. brasiliensis* (*4*). Subsequent molecular, phylogenetic, *in silico* DNA-DNA hybridization and average nucleotide identity (ANI) analysis led to the elevation of this taxon to the species level (*5*-*8*).

*Pectobacterium* species are known to co-infect (*9*-*11*) and show all three inter-species interaction behaviors -competition, cooperation and commensalism (*12*). Until 2013, *Dickeya* spp. and *P. atrosepticum* were dominant in western European countries, like the Netherlands and Switzerland. After the introduction of *P. brasiliense* in western Europe in 2013, its abundance increased rapidly and it is now detected in almost 70-80% of the blackleg diseased plants (*13–15*). Strain specificity towards infecting certain host plants (*3*) and the ability to cause latent infections (*16*) are characteristics of *P. brasiliense*. This species shows regular emergence of new lineages, also evident from the observed high heterogeneity and a broad range intra-species ANI (*17*). Recent pangenome studies on *Pectobacterium* genus have shown a growing pangenome indicating that not all diversity is captured in this genus yet (*17*, *18*).

In prokaryotes, various mechanisms are responsible for the evolution of such heterogeneous strains, such as mutations and horizontal gene transfer (HGT), for example via plasmids and bacteriophages (short phages). Genomes for many species belonging to the *Pectobacterium* genus were assembled at single contig level with minimal evidence of autonomous plasmids (*19*-*21*) and yet, computational analyses predicted the presence of plasmids in other 89 accessions (*22*). Some phages can integrate into a prokaryotic genome through the lysogenic pathway and are called prophages. A study on a small collection of 54 *Pectobacterium* spp. genomes showed the presence of prophage-like sequences (*23*). These prophages can influence physiological functions and provide fitness advantages to the host bacteria by mechanisms such as superinfection exclusion and harboring virulence factors and antibiotic resistance genes (*24–28*), also noted in *Pectobacterium* pathogens (*29*). From a wider perspective of population expansion, prophages are known to contribute to the competitive success and dispersal of a prokaryotic strain from the environmental reservoir (*30*, *31*). Therefore, it is critical to understand the role of prophages in species and strain diversity, and its contribution to population structure, evolution and epidemiology.

Two distinct groups of *P. brasiliense* strains were detected during annual surveys from 2016 to 2018 in the Netherlands: a clonal lineage with a high ability to cause BL and another, relatively diverse group unable to cause BL in potatoes in field trials (*17*). A subsequent comparative pangenomic analysis enabled the design of new functional assays to detect these BL-causing isolates (*32*). However, a new group of BL-causing isolates that escaped these assays were detected during ongoing annual surveys. A common observation is the emergence of new lineages of *P. brasiliense* in the Netherlands followed by their dominance over the existing lineages. We therefore extended the *Pectobacterium* genus pangenome by combining a genome collection from local epidemiological surveillance with global data to understand the mechanism of appearance and succession of the *P. brasiliense* strains. Further, we investigated the contribution of mobile genetic elements, and prophages in particular, in the growing *Pectobacterium* genus pangenome. Our results showcase an application of the pangenome approach to trace such mobile genetic elements, delineate their dynamics and provide mechanistic insights into pangenome growth.

## Results

### Emergence of blackleg-causing *P. brasiliense* isolates that evade diagnostics

Yearly inspection, coupled with PCR-based identification of plant pathogens, indicates that after 2013 *P. brasiliense* has emerged as the dominant pathogen in the Netherlands, replacing *Dickeya* spp. In recent years, we have observed a higher prevalence of *P. brasiliense* on blackleg and softrot infected plants, wherein 95% of symptomatic plant material show infection by *P. brasiliense* (Figure 1a). From 2018 to 2020, *P. brasiliense* isolates were sampled from symptomatic and asymptomatic potato (*Solanum tuberosum*) plants in the Netherlands. These isolates were genotyped as BL-causing or BL-non-causing based on diagnostic quantitative PCR assays (*32*). Subsequent field trials showed that some of the presumed non-causing *P. brasiliense* isolates did induce the blackleg phenotype, implying false negative PCR diagnosis. These isolates are henceforth referred to as ‘FN-Pbr’.

**Figure 1:**
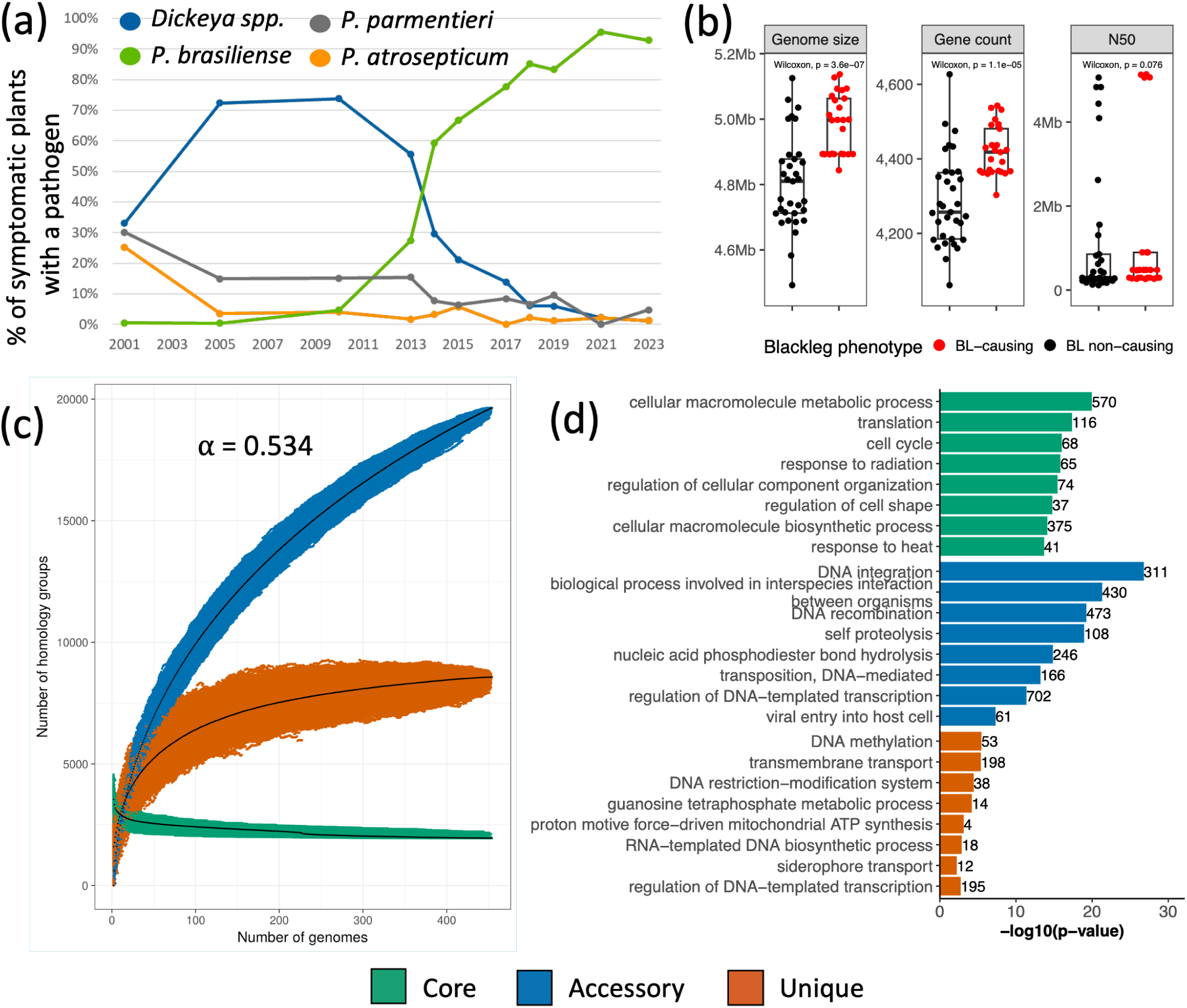
(a) Percentage of different pathogens identified from the plants showing BL symptoms during the annual inspection in the Netherlands. (b) In the Dutch collection, BL-causing isolates have larger genomes and more protein-coding genes than BL-negative isolates. (c) The *Pectobacterium* pangenome has a stable core but increasing accessory genome. Heaps’ law *alpha* = 0.534 is *<* 1 and implies an open pangenome. (d) Functional enrichment of Gene Ontology biological processes in the core, accessory and unique pangenome.

We also detected multiple *Pectobacterium* species on the same infected plant material during the inspection in the Netherlands. To assess the relation between the newly identified FN-Pbr isolates and the existing BL-causing isolates and to understand potential inter-species interactions, we sequenced 58 *Pectobacterium* spp. isolates collected in the Netherlands. This collection included 28 *P. brasiliense* isolates, 10 of which were FN-Pbr. Genome assemblies from public databases were leveraged to study FN-Pbr evolution at high resolution and further capture the genomic diversity in *Pectobacterium* genus. At the time of analysis, the NCBI database contained 450 *Pectobacterium* spp. genomes. We added 396 high-quality genomes of these after curation (see Methods), resulting in a collection of 454 genomes representing 22 *Pectobacterium* species (Supplementary Table S1, Supplementary Figure S1).

### A growing *Pectobacterium* genus pangenome

To study the overall sequence diversity in our collection of genomes, we first performed pairwise ANI comparisons. Within the *P. brasiliense* clade, we observed a broad range of intra-species ANI scores: 9.8% of *P. brasiliense* genome pairs had ANI scores of 96% or less, 1.8% of pairs were with scores of 95% or less. Additionally, a genus level ANI comparison showed at least five distinct *Pectobacterium* species for which inter-species ANIs were closer to 95%, a commonly used species delineation cutoff (*33*, *34*) (Supplementary Figure S2). For example, a clade with *P. versatile*, *P. carotovorum* and *P. odoriferum*; and another clade with *P. polaris* and *P. parvum*. Together, these results demonstrate a high genetic variability in *Pectobacterium* species, in particular for *P. brasiliense* in the Netherlands. Additionally, BL-causing *P. brasiliense* isolates had significantly larger genomes (Wilcoxon-test, *p* = 3.6 *×* 10*^−^*^7^) and more protein-coding genes (*p* = 1.1 *×* 10*^−^*^5^) than non-causing ones (Figure 1b). This was not due to differences in assembly quality, as there was no significant difference (*p* = 0.076) in median assembly N50 values between the two groups and all genomes had BUSCO completeness *≥* 99%.

For a reference-free comparison of this heterogeneous genus, we constructed a genus-level pangenome of 454 *Pectobacterium* spp. genomes using PanTools (*35*). In our pangenome 1,977,865 protein-coding genes were clustered into 30,156 homology groups, which were further categorized into 1,949 core (present in all genomes), 8,571 unique (exclusive to individual genomes) and 19,642 accessory groups. Fitting a Heaps’ law model to homology group counts as a function of newly added genomes provides an estimate of the pangenome openness, where a decay rate *α <* 1 implies an open pangenome and *α >* 1 a closed one (*36*). In line with the earlier, smaller pangenomes (*17*,*18*), the *Pectobacterium* spp. pangenome is still open, with a decay rate *α* of 0.534 (Figure 1c). In the species-level sub-pangenomes extracted from the genus-level pangenome, *P. versatile* and *P. brasiliense* are represented by significantly more genomes than others but surprisingly had the lowest decay rates of 0.501 and 0.538 respectively (Supplementary Figure S3, Supplementary Table S2). After more than doubling the number of genomes in the genus-level pangenome compared to the previous version, the core remained stable with a small decrease of 83 homology groups (Supplementary Figure S4). The increase in the overall pangenome size by 7,815 homology groups was mainly attributed to the accessory category, with 6,474 additional homology groups. This increase was mostly contributed by *P. brasiliense* and *P. versatile*, with 7,290 and 6,384 accessory homology groups respectively (approx. 50% of the homology groups in the respective sub-pangenomes) (Supplementary Table S2).

### A new blackleg-causing *P. brasiliense* lineage

A maximum likelihood phylogenetic tree based on the alignment of 1,949 core genes revealed that all 25 BL-causing *P. brasiliense* isolates were part of a monophyletic clade of 48 isolates (Figure 2). This BL-causing clade was split into two sub-clades: one of 35 isolates enriched for TP-Pbr, the other of 13 isolates enriched for FN-Pbr. Intra-clade pairwise comparison of the isolates within these sub-clades showed 100% ANI. Based on the core phylogeny and ANI comparison, it is evident that FN-Pbr and TP-Pbr represent clonal lineages radiating from a BL-causing *P. brasiliense* common ancestor Supplementary Figure S2). Only one strain (NAK399) from the FN-Pbr sub-clade did not cause any BL phenotype in the field trials (0.5% infected plants).

**Figure 2:**
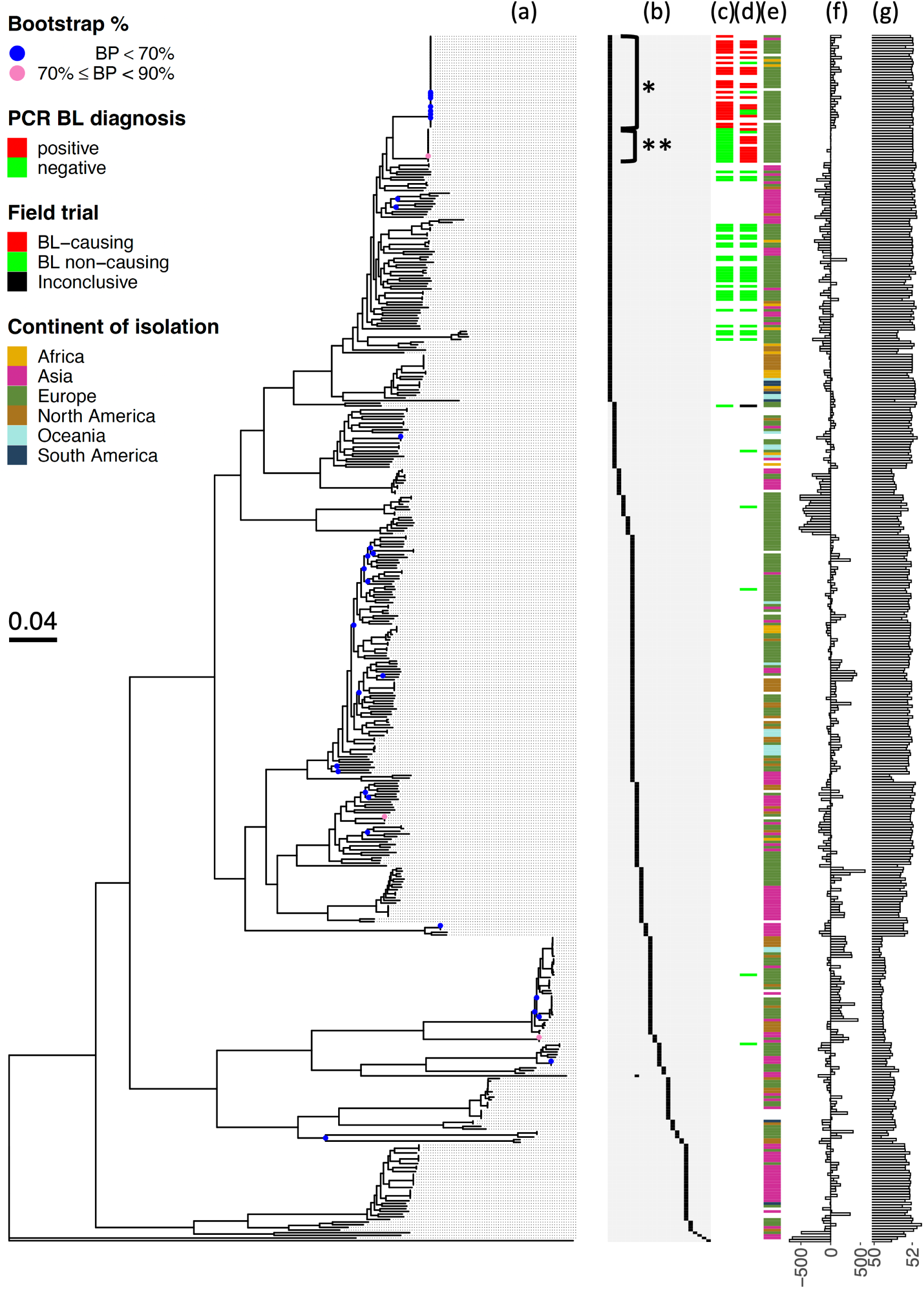
(a) A robust maximum likelihood tree generated using 1,949 core gene alignments. Nodes with bootstrap values *<* 90% and *<* 70% are colored pink and blue respectively. Metadata (from left to right): a species key (b), where each column denotes a species ordered from left to right: *P. brasiliense*, *P. polaris*, *P. parvum*, *P. quasiaquaticum*, *P. aquaticum*, *P. versatile*, *P. carotovorum*, *P. odoriferum*, *P. actinidiae*, *P. parmentieri*, *P. wasabiae*, *P. punjabense*, *P. polonicum*, *P. atrosepticum*, *P. peruviense*, *P. zantedeschiae*, *P. betavasculorum*, *P. aroidearum*, *P. carotovorum subsp. carotovorum*, *Pectobacterium* sp. CFBP8739, *P. colocasium*, *P. fontis* and *P. cacticida*. *: Clade enriched for TP-Pbr isolates. **: Clade enriched for FN-Pbr isolates. PCR based diagnosis of BL-causing ability of the isolates (c), BL phenotype during field trials (d), continent on which the isolate was collected (e), difference in number of protein-coding genes with respect to the median of 4,363 genes across *Pectobacterium* genus (f), and GC% (lower limit of the bar plot shown is 50%) (g).

To gain broad functional insights of our pangenome, we performed a gene ontology enrichment analysis on core, accessory and unique gene sets. The *Pectobacterium* core genome was enriched for elementary processes such as cell cycle, translation, heat response, macromolecule metabolism, ribosome assembly, etc. The unique genome was enriched in DNA methylation, siderophore transport and lipid transmembrane transport. The accessory genome was enriched for DNA integration, recombination, transposition, viral entry into host, inter-species interaction, etc. (Figure 1d, Supplementary Table S3).

### Contribution of prophage-like elements in the *Pectobacterium* pangenome

Intrigued by our observation of the heterogeneity in *P. brasiliense* and enrichment of biological processes related to mobile genetic elements, we hypothesized such elements as an underlying factor in pangenome growth. Therefore, we systematically analyzed mobile genetic elements in *Pectobacterium* genomes using the geNomad pipeline (*37*) (see Methods for details). A total of 1,369 prophage-like regions were identified in the 454 genomes, where all but one genome contained at least one prophage. Cumulatively, genes from prophages accounted for 4,801 (15.9%) of the pangenome homology groups. Among these prophage homology groups, 3,286 (16.7%) were accessory, 1,477 (17.2%) unique and only 38 belonged to the core genes in *Pectobacterium* genus pangenome (Figure 3a). Similarly, predicted plasmids were represented by 10,544 homology groups. A comparison of clonal isolate genomes (*ANI >* 99.9%) from the TP-Pbr clade (*n* = 35) showed that geNomad predicted plasmids in fragmented genome assemblies (*n* = 26) but not in the single chromosome assemblies (*n* = 9). Furthermore, the plasmid homology groups from the isolates with fragmented genome assemblies could be mapped in tandem onto the chromosome-level assembly, suggesting a false-positive identification. Alternatively, these can be integrated plasmids in the single chromosome genomes, also known as episomes; this is supported by the consistent observation of contigs of length 293kb and 89kb as putative plasmids in the fragmented genome assemblies. Nevertheless, this renders the plasmid prediction results unreliable.

**Figure 3:**
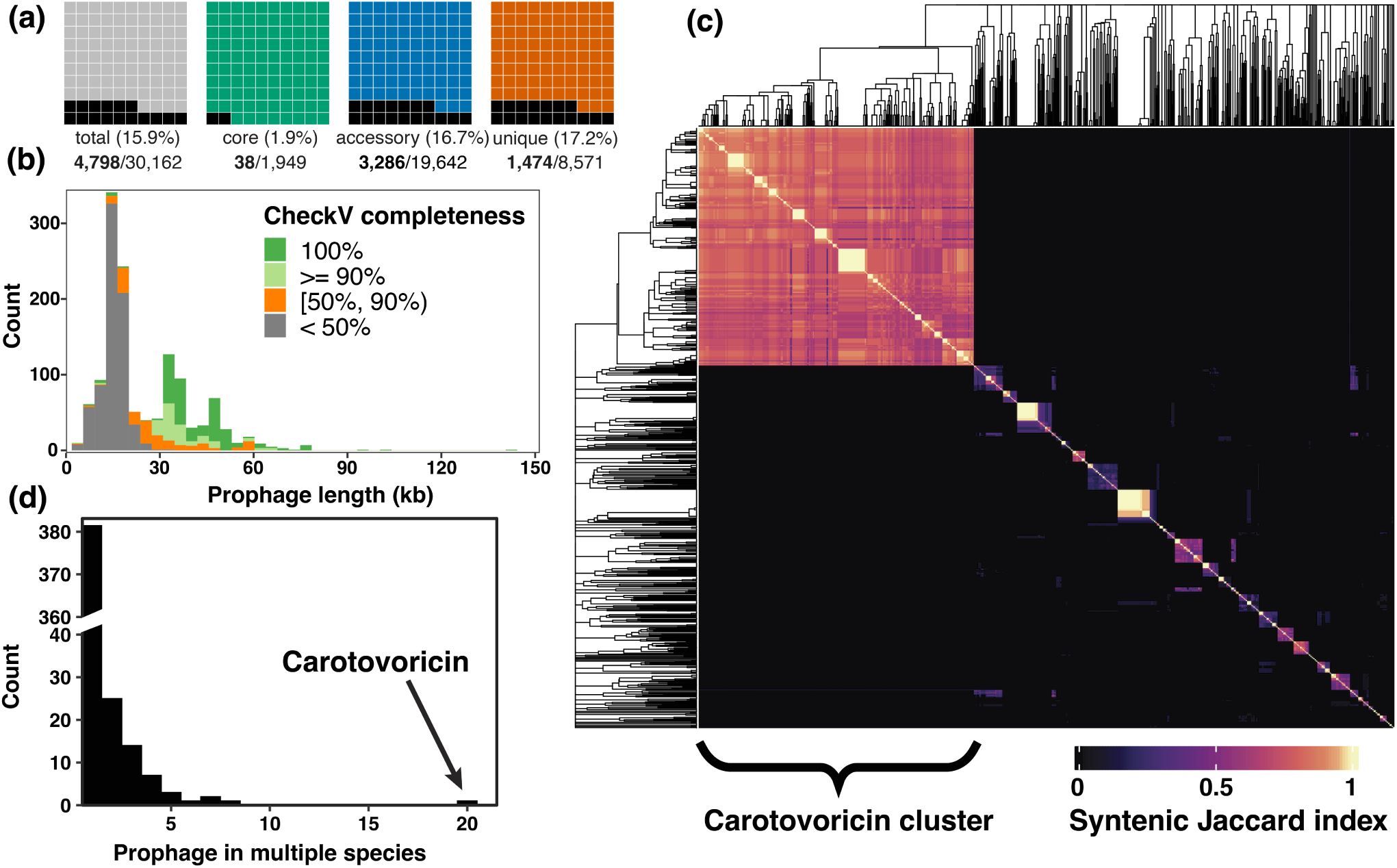
Prophage analysis. (a) Fraction of pangenome homology groups corresponding to prophages. (b) Prophage length distribution. (c) Clustering based on homology group content, represented by a heatmap of syntenic Jaccard index values. The largest cluster of carotovoricin (CTV) is annotated in the heatmap. Complete linkage dendrogram was cut at the node height of CTV cluster to identify groups of orthologous prophages. (d) A histogram showing prophage backbone present in one or multiple *Pectobacterium* species. The CTV, a prophage-like element present in 20 species is marked with an arrow.

We used CheckV (*38*) to assess the completeness of the putative prophages and found 731 prophage-like elements with a completeness score of 50% or lower and 423 prophages with 90% or higher completeness (Figure 3b, Supplementary Table S5). The shortest prophage with 95% or more completeness was 5.4kb long. The prophage length distribution shows three distinct peaks, with the highest peak at 16kb mostly representing incomplete prophages; a peak at 35kb representing prophages with completeness *>* 90%; and a peak at 50kb comprising complete prophages. At the species level, *P. brasiliense* and *P. versatile* prophages contributed to *∼* 14% of their respective pangenomes (Supplementary Table S4). This distribution is similar to the prophage fraction in the accessory genome of other highly dynamic bacterial pangenomes like *E. coli* and *S. enterica* (*39*). Since the BL-causing *P. brasiliense* isolates have larger genomes, we compared the total prophage length between the two groups for a potential explanation. We observed that the BL-causing isolates had significantly larger median total prophage length (*t*-test, *p* = 1.1 *×* 10*^−^*^4^). This 50kb difference observed in total prophage length accounts for 25% of the observed 200kb disparity in median genome sizes between these two categories (Supplementary Figure S5).

The tailed bacteriophage class *Caudoviricetes*, with a double stranded DNA genome, was represented most abundantly (*n* = 1, 357) in the *Pectobacterium* pangenome. Although much less abundant, other classes of viruses included filamentous phages of the *Inoviridae* (*n* = 8) and *Microviridae* (*n* = 2) families, a virus from *Patatavirales* order (*n* = 1) and a 5.2kb unclassified prophage (Supplementary Figure S6). *Inoviridae* and *Microviridae* are non-enveloped single-stranded DNA viruses, whereas *Patatavirales* is the largest order of non-enveloped positive-strand RNA viruses of plants and contains a single family *Potyviridae*, accounting for 30% of all known plant viruses (*40*). While it was interesting to identify a plant virus of the order *Patatavirales* in the *P. parvum* genome assembly GCF 011378945.1, caution must be taken, as phages detected on independent contigs are difficult to ascertain as true integrations. Among the less abundant phages, host DNA integration evidence was observed for *Inoviridae* family prophages (Supplementary Table S5), thus these can be considered true prophages. The *Microviridae* and *Patatavirales* prophages were detected on an independent contig without flanking host genes and are likely the result of lytic viruses or sample contamination.

### Generalist prophages of *Pectobacterium* species

Bacteriophages have high genetic diversity and mosaicism is a hallmark of their genomes, primarily driven by horizontal gene transfer (*41–44*). Given the variation in prophage length and the dominance of a single bacteriophage class, finding orthologous prophages would help in the characterization of prophage diversity and dynamics in the *Pectobacterium* pangenome. Therefore, we compared putative prophages by calculating syntenic Jaccard indices of the prophage homology group signatures. CTV, a conserved 16kb phage tail-like bacteriocin occurring in 428 genomes was used to decide the prophage grouping threshold (Figure 3c) (see Methods). As a result, we identified 436 clusters of orthologous prophages (named phage grp 1 to phage grp 436) and their respective cluster representative (Supplementary Figure S7, Supplementary Table S6). These representative prophages covered a total of 4,493 homology groups (94% of the total 4,801), implying minimal loss of information during clustering.

Most phages are species-specific and even strain/serovar/pathovar specific (*45–47*), though broad range phages targeting different species also exist (*48*, *49*). To identify such generalist phages infecting multiple *Pectobacterium* species, we checked prophage clusters for the presence of multiple species. We identified 54 generalists and 382 species specialists (Figure 3d). For example, clusters phage grp 71 (Figure 4a) and phage grp 36 with 24 and 21 prophages respectively, included complete prophages targeting multiple *Pectobacterium* species and their hosts were collected from a wide spatiotemporal context (Supplementary Table S6). No association was found between the *Pectobacterium* isolates harboring phage grp 71 phages and plant infection, as isolates were collected from both symptomatic and asymptomatic plants. Furthermore, none of the isolates harboring phage grp 36 phages were sampled from infected plants. *P. versatile* was the most frequent host of the generalist prophages, targeted by 34 out of 54 prophages (Figure 4b-c). The diverse ecological niches of *P. versatile* isolates, spanning water, soil and infected plants, likely contributed to their susceptibility to generalist phages. In contrast, the closely related species *P. aquaticum* and *P. quasiaquaticum* collected from freshwater bodies in France did not share any prophages, yet did share prophage signatures with other *Pectobacterium* species.

**Figure 4:**
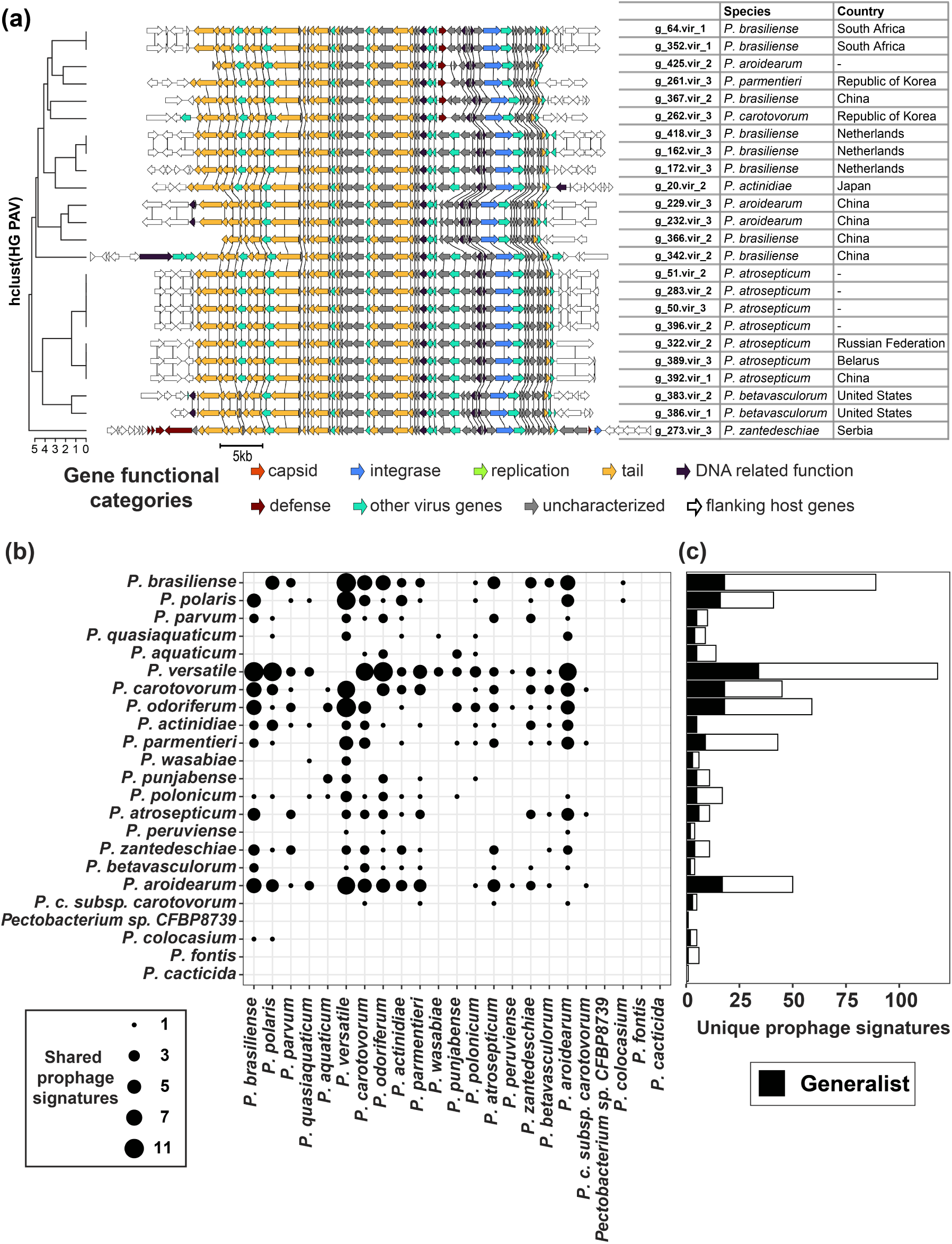
(a) Prophage signatures of 22 members of the phage grp 71 cluster. Flanking genes are colored white and the genes belonging to the same homology groups are connected with vertical lines. (b) Generalist prophages shown as the overlap of prophage signatures between *Pectobacterium*species pairs. (c) Number of unique prophage signatures per species. The fraction of generalist prophages is indicated in black.

### Prophage dynamics in the BL-causing clonal lineages

We further explored the prophages of isolates in the BL-causing clade, specifically to understand the intra-lineage variation and horizontal transfer with other *Pectobacterium* species. We detected four clusters with intact prophages exclusive to the BL-causing clade (Figure 5a). First, a 49kb intact prophage was found in all but two isolates from this clade (phage grp 45 in Supplementary Table S6). This prophage carried a YdaS/YdaT toxin-antitoxin (TA) system and a DNA methyltransferase gene, important elements of bacterial defense systems (Figure 5b). The same flanking genes are found across all genomes, which suggests a single insertion event of this prophage in the common ancestor of TP-Pbr and FN-Pbr isolates followed by vertical transmission.

**Figure 5:**
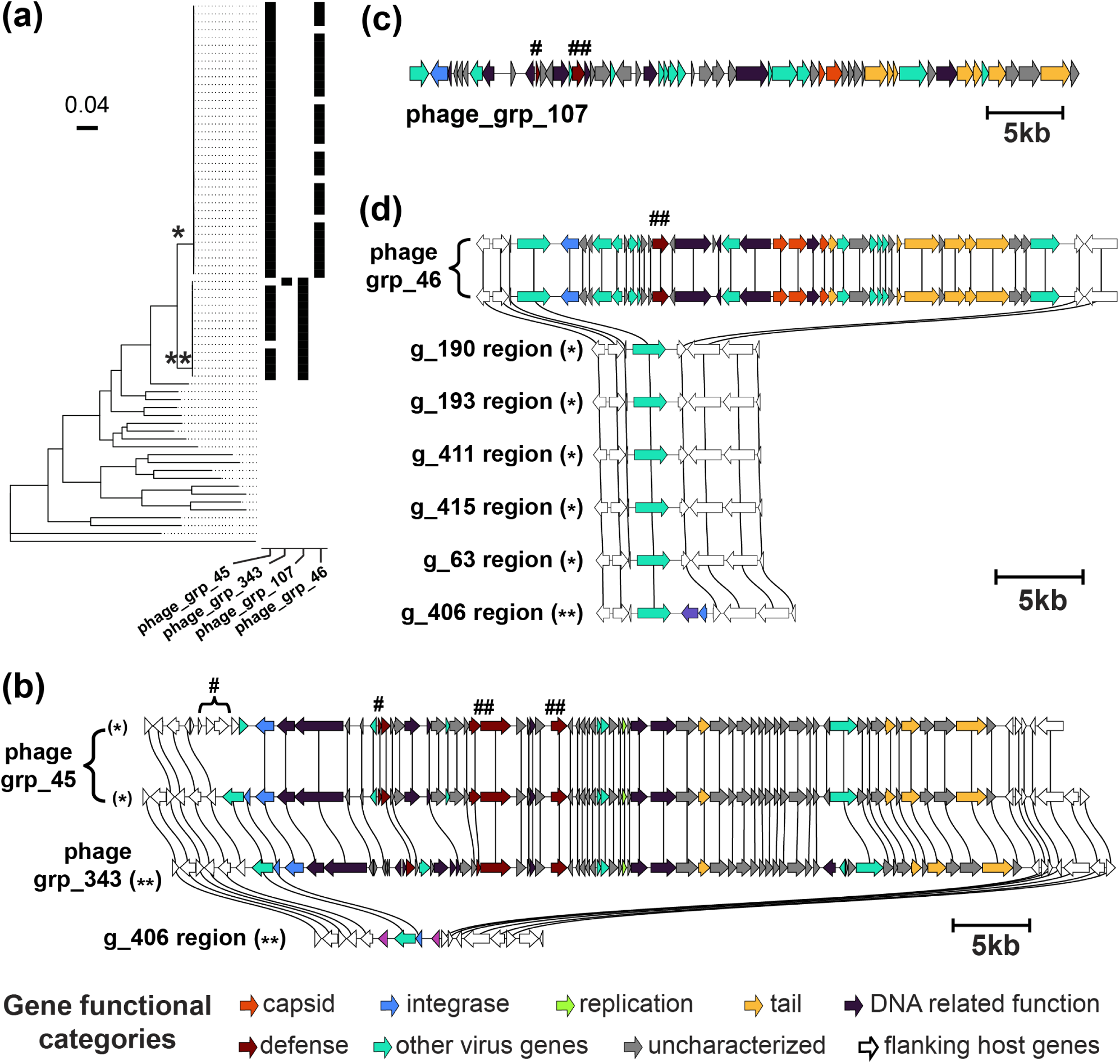
(a) Presence-absence (black-white) of the four prophages exclusive to BL-causing *P. brasiliense* lineage are shown next to a subtree for this clade. *: TP-Pbr clade, **: FN-Pbr clade. The remaining nodes in the tree belong to one representative *Pectobacterium* species. (b) BL-causing clade-specific prophage homology group signatures: four regions showing two representative prophages from the orthologous phage group phage grp 45, a single prophage from genome g 400 belonging to phage grp 343 group, and a region from genome g 406 in which the prophage is absent. (c) FN-Pbr clade-specific prophage homology group signature: a representative from the orthologous phage group phage grp 107. (d) TP-Pbr clade-specific prophage homology group signatures comparison: two representative prophages from phage grp 46 group and regions from the five TP-Pbr genomes and the single FN-Pbr genome in which this prophage is absent are shown. For all figures, flanking genes are colored white and genes in the same homology groups are connected with vertical lines. Bacterial defense systems are marked as #: TA-system and ##: methyltransferase. BL diagnosis is marked as *: TP-Pbr, **: FN-Pbr.

A single FN-Pbr clade isolate sampled in 2019 in the Netherlands showed an insertion of a 50kb prophage (phage grp 343 in Supplementary Table S6). Interestingly, this prophage had a very similar homology group signature to phage grp 45 prophages, with some rearrangements, and was integrated at the same site in the host genome (Figure 5b), implying a mosaic relationship. These prophages could potentially be clustered together when a relaxed threshold is applied. Nevertheless, they are distinct in gene content and structure and our approach effectively captures phage mosaicism and dynamics across multiple genomes. Another 44kb prophage was detected in all of the FN-Pbr isolates (phage grp 107 in Supplementary Table S6, Figure 5c). In fragmented genome assemblies of seven FN-Pbr isolates, this prophage was split over two or more contigs. However, our pangenomic homology group signature matching allowed merging these fragments into a single prophage signature, demonstrating the use of a pangenomic approach in studying mobile genetic element dynamics. Interestingly, this prophage harbored a methyltransferase and antitoxin gene of the TA-system, although the toxin component was missing.

Finally, a 31kb prophage was detected exclusively in 30 TP-Pbr isolates collected between 2012 and 2020, mainly in the Netherlands (phage grp 46 in Supplementary Table S6). The five *P. brasiliense* isolates in this clade lacking this prophage were sampled in 2016 or later in Belarus and the Netherlands (Figure 5a,d). Prophage signature comparison shows a remnant region in FN-Pbr genomes (g 406). Therefore, an insertion in the common ancestor of TP-Pbr and FN-Pbr clades and two independent excision events, first in the FN-Pbr clade and another in the five isolates of TP-Pbr clades likely explains these results. In summary, we captured ongoing prophage and immune system dynamics in the clonal isolates based on the robust core phylogeny.

## Discussion

During annual sampling of soft-rot *Pectobacteriaceae* in the Netherlands, we detected BL-causing *P. brasiliense* isolates that escaped the previously designed BL-phenotype assays (*32*). To provide a broader context for the emergence of this isolate, we applied a pangenome approach to the *Pectobacterium* genus. Compared to the previous pangenome, we observed a stable core but an almost 50% increase in the acces-sory genome. Prophages contributed significantly to the *Pectobacterium* genus pangenome, mostly non-core genes. Further characterization of these prophages revealed the presence of both generalist and specialist prophages of *Pectobacterium* species.

The genetic diversity of *P. brasiliense* is evident from the broad range of intra-species ANI scores (94-99%). Pathogens can evade diagnosis by natural selection of isolates that acquire mutations in assay target genes. Diagnostic assay escape is not new for soft-rot pathogens. Previously, the Dia-A primer developed to detect *D. dianthicola* failed to detect a *Dickeya* outbreak in the USA because of target gene deletion (*2*). In this and our case, the newly evolved isolates were very similar to those for which assays were developed (for *P. brasiliense*, 97.7% ANI). This highlights the challenges in designing stable phenotype diagnosis assays for soft-rot pathogens like *P. brasiliense* with high genomic diversity and an open pangenome. To address this, we propose monitoring as described in this study and periodic pangenome updates.

Bacterial species show a wide range of prophage dissemination, from 50% of the genomes in actinobacterial species harboring one or more prophage-like elements (*50*) to 99.5% of *Acinetobacter baumanii* genomes (*51*). Higher dissemination of prophages is known to drive the plasticity of bacterial genomes and con-tributes to pathogenicity and virulence (*51*). In our collection, including *Pectobacterium* spp. isolates from all over the world, 99.7% of the genomes had one or more prophage-like elements, implying high genome plasticity at both genus and species levels. We further showed that prophages account for *∼* 16% of the *Pectobacterium* genus pangenome, resulting in an open pangenome. We observed a significant difference in mean genome size between the BL-causing and non-causing *P. brasiliense* isolates. Prophages explain a quarter of this observed genome size difference. Bacteria and their phage predators are locked in an arms race and it is possible that to counter these phages, prey bacteria are under selection pressure to acquire additional mobile genetic elements such as plasmids, integrative conjugating elements and insertion sequences. We did not find convincing evidence for the presence of plasmids in *Pectobacterium* species as we observed a discrepancy in plasmid prediction on high quality versus fragmented genome assemblies. It remains to be seen whether other mobile genetic elements exist and how they interplay with the prophages in *Pectobacterium* species.

As observed in *P. brasiliense* lineages from our collection, lineage-specific prophages are also detected in other pathogens, for example Φ*RS*551 in *Ralstonia solanacearum* (*52*) and Φ*RE*2010 in *Salmonella enterica* (*53*). Genes in such prophages are known to be transcriptionally active and up-regulated post-infection in different plant pathogens, for example in *P. brasiliense* (*54*) and *R. solanacearum* (*55*, *56*). These prophages modulate the mobility and virulence of bacterial pathogens and may provide a compet-itive advantage over other closely related species not bearing prophages (*29*, *54*). Similar examples of horizontally acquired mobile elements modulating virulence and quorum sensing are also reported for the *Pectobacterium* species (*29*, *57*, *58*). Moreover, annual sampling across the Netherlands enabled us to capture ongoing gain-loss dynamics of prophages, even within clonal lineages, and therefore highlights the contribution of prophages to genome plasticity. These lineage-specific prophages in *P. brasiliense* probably provide a competitive advantage over other closely related species and the active dynamics of prophages enable such pathogens to continuously radiate novel lineages as observed.

We also showed the existence of generalist prophages targeting multiple *Pectobacterium* species. Such prophage identification across species validates host recognition by the phage, an important characteristic for determining phage host range. Phage-host interaction networks often show modularity (*49*, *59*). Taxo-nomic separation of hosts due to structural and metabolic differences is the intrinsic driver of such modular host ranges (*45*), emphasizing the need to include the host phylogeny while analyzing phage host ranges. A pangenome approach to trace prophages shown here allows such paired study of host range and host phylogeny estimated from the core genome. These phage host range studies have important applications such as phage typing, controlling microbial community composition and phage therapy.

Biocontrol of plant pathogens using phages provides an environmentally friendly alternative to antibiotics (*60*). Although lytic phages are more suitable phage therapy candidates, studying the dynamics of lysogenic phages will enable learning of the host-detection mechanisms employed by phages in general. Recently, several studies have shown the effectiveness of this for *Pectobacterium* species and other plant pathogens (*61–67*). However, just as antibiotic resistance, the development of phage resistance in bacterial hosts (*68*) is a major bottleneck in phage therapy applications. Prophages also carry various defense systems, such as TA-systems, restriction-modification system and CRISPR (*69*, *70*), that can provide immunity to the host bacteria against both lytic and lysogenic phages (*71*). All of the BL-causing *P. brasiliense* exclusive prophages harbored at least one defense system. Therefore, a comprehensive characterization of prophage diversity and its interplay with bacterial defense systems can pave the way for smarter development of phage therapy applications. Additionally, a broader pangenome study that includes pathogens sharing niches with *Pectobacterium* species could help to discover prophages with cross-genus range.

However, such large-scale comparative analysis to study prophage dynamics comes with several challenges, such as reference bias and prophage overestimation because of fragmented assemblies (*72*). Our approach overcomes these by combining prophage identification with the graph pangenome approach to merge fragmented prophages and correctly estimate their abundance in pangenome. We further used homology groups to identify orthologous prophages and constructed a non-redundant prophage set for *Pectobacterium* species. Thus, our approach provides a novel and efficient strategy to screen and select bacteriophage candidates, also for developing phage therapy applications.

In conclusion, our study demonstrates an important contribution of prophages in the growing *Pectobacterium* species pangenome. We identified an ongoing bacteriophage-mediated exchange of genes in the new lineage of *P. brasiliense* as well as highly conserved prophage-like elements in the *P. brasiliense* genomes. Our results thereby emphasize the importance of routine sampling and pangenome analysis in understanding the evolutionary trajectories of bacterial plant pathogens.

## Methods

### Sample collection and phenotyping

During the annual screening and field inspection conducted by the Dutch General Inspection Service for agricultural seeds and seed potatoes (NAK) in the Netherlands from 2018 to 2020, *Pectobacterium* spp. strains were collected from symptomatic potato plants and non-symptomatic potato tubers. Non-symptomatic potato peels were crushed in water and incubated in pectate buffer before spreading on single-layer crystal violet pectate (CVP) plates. Characteristic cavity-forming colonies were sub-cultured on the CVP and nutrient agar plates to obtain pure strains and stored at *−*80*^◦^*C in 15% glycerol with half-strength nutrient broth.

BL-causing ability of the *P. brasiliense* isolates was tested on potato cultivar Agria during 2019 to 2023 field trials. Isolates were grown on nutrient agar plates for one day at 28*^◦^*C. Suspensions of OD_600_ = 0.1 were prepared in 10 mM phosphate buffer (pH 7.2) and diluted an extra 100*×*, resulting in a suspension of about 106 cfu/ml. Tubers were submerged in the solution and brought under a *−*0.07Pa vacuum. The vacuum was kept for 10 minutes, after which tubers were left submerged for another 15 minutes. Tubers were air-dried before planting and the ability to cause BL was scored as the development of typical plant symptoms (*14*). Isolates that induced BL phenotype in *≥* 30% plants were categorized as BL-causing, in *<* 5% plants as non-causing and the remaining were labelled inconclusive.

Additionally, 35 *Pectobacterium* species isolates were included from the Netherlands Institute for Vectors, Invasive plants and Plant health (NIVIP) collection.

### Genome sequencing, quality control and annotation

Whole genome sequencing and genome assembly were performed for 23 *P. brasiliense* isolates from the NAK collection and 35 isolates from the NIVIP collection as described in Jonkheer et al. (*17*) and Blom et al. (*73*) respectively. Publicly available genomes from *Pectobacterium* species and related BioSample metadata were downloaded (date: 29/08/2022) using the NCBI E-utility tool (*74*). For uniform taxonomy validation of our collection, we combined in-house and NCBI type-strains to perform an independent ANI comparison and taxonomy correction. Of the 508 genomes, 26 marked as “anomalous”, “excluded” or “replaced” by NCBI were excluded. We used BUSCO (v5.2.2, database: “enterobacterales odb10”) protein to evaluate genome assembly completeness and removed 21 genomes that had less than 99% “complete BUSCOs” (*75*). Finally, seven genomes were found to be duplicated and hence only one copy was kept for further analysis. The remaining 454 genomes were annotated using Prokka (v1.14.6) (*76*). InterProScan (v5.56-89.0) was used to annotate genomes with protein family and domain information from the InterPro database (*77*, *78*). EggNOG-mapper (v2.1.10) was used to annotate proteins by transferring annotations from the EggNOG database (v5.0.2) (*79*, *80*).

### Pangenome construction and analysis

A *Pectobacterium* spp. pangenome was built using PanTools (v4.1.1) (*35*, *81*). Both InterProScan and EggNOG annotations were added to the PanTools pangenome database using the *add annotations* module. To group genes into homology groups, PanTools’ *optimal grouping* command tests eight different similarity cutoffs and evaluates the grouping of single-copy BUSCO genes. The time required to optimize the grouping increases quadratically with the number of genomes. To efficiently determine the relaxation setting for grouping of genes, we used two subsampling approaches. First, genomes of the type strain for each species were used. In the second, the *k*-mer tree built using *kmer classification*command was cut into 20 clusters (number of *Pectobacterium* species) and a genome was selected from each cluster at random. This second method of random subsampling was repeated 100*×* to generate 100 subsets of *Pectobacterium* genomes. The optimal grouping step of PanTools was performed on these subsets and the relaxation setting with lowest average *F* -score was selected. This resulted in a relaxation setting of four, which corresponds to 65% sequence similarity, 0.05 intersection rate for the *k*-mer set comparison, a 7.2 inflation rate for Markov clustering and a contrast factor of five.

### Phylogenetic analysis

Core genes were used to construct a maximum likelihood phylogenetic tree using the PanTools *core phylogeny* command (*82*). Phylogenetic trees were processed using R (v4.2.1) packages Ape (v5.7.1) (*83*, *84*) and treeio (v1.22.0) (*85*) and data was visualized with R packages ggtree (v3.6.2) (*86*) and ComplexHeatmap (v2.15.1) (*87*).

### Prophage detection and quality control

Prophages were detected in genomes using the geNomad (v1.5.0) pipeline and checkV (v1.0.1) was used for virus genome completeness assessment (*38*, *88*). Host sequence regions at the prophage boundaries predicted by CheckV were trimmed to remove the contamination resulting from host genes. 139 prophages without any gene of viral origin were excluded from downstream analysis. In fragmented genome assemblies, prophages may be split over multiple contigs, leading to an over-estimation of the number of prophages.

We used homology group information of the orthologous prophages from the high quality genome assem-blies to identify such fragmented prophages (*n* = 145) and merged them into 61 prophages. Merging was considered confident if the merged prophages together covered at least 80% of the closest matching orthol-ogous prophages (*n* = 50 from 110 fragmented prophages) and the rest were excluded (*n* = 11). Finally, the shortest prophage detected with *>* 90% completeness was of length 5.2kb and hence prophages smaller than 5kb were also excluded, resulting in 1,369 prophages *≥* 5*kb* in length.

### Orthologous prophage detection

Each putative prophage region was represented as a sequence of homology groups from the pangenome, de-fined as the prophage signature. Prophage similarity was quantified by evaluating the syntenic conservation of homology groups between two prophage signatures. To quantify such syntenic conservation, we applied dynamic programming to align two prophage signatures and calculate the pairwise Jaccard index. Only syntenicly matching homology groups between a pair of prophage signatures are considered for determining the size of the intersection for the Jaccard index calculation. During dynamic programming scoring, a score of 5 was used for a match and -2 for a mismatch of homology groups between two prophage signatures being compared. To account for the natural variation in prophages due to the mosaicism and evolution, a maxi-mum gap length of two was allowed. A syntenic Jaccard index was considered significant if two prophages had the longest local subsequence with a chain length, i.e. the number of syntenicly matching homology groups, of at least five. An all-vs-all syntenic Jaccard index matrix was clustered using the complete linkage hierarchical clustering function ‘hclust’ in R. A conserved phage tail-like bacteriocin, CTV, was present in 428/454 *Pectobacterium* genomes and clustered together. We used CTV’s intra-cluster similarity as a threshold to decide the grouping of similar prophage-like elements into clusters. The complete-linkage dendrogram was cut at height 0.66, which was just above the node height for the CTV cluster, resulting in 436 prophage clusters. Here, the complete-linkage clustering ensures that each resulting cluster includes prophages with intra-cluster similarity equal to or higher than the chosen cut-height or similarity thresh-old. A cluster representative backbone was selected using criteria in the following order: highest checkV completeness score, longest prophage length and highest average intra-cluster Jaccard index.

### Prophage cluster visualization

A 5kb flanking region for each prophage was included to provide a broader context to understand integration at the same or different sites in the genome. An R script was written to extract the homology groups for prophages in a cluster and the data was stored in required JSON format by clustermap.js (https://github.com/gamcil/clustermap.js). This JSON data was used in clinker (*89*) to generate interactive plots to visualize orthologous prophages. Prophage genes were manually classified into broad functional categories based on the PFAM and COG description: integrase, capsid, replication, defense, DNA-related functions, phage tail-related, other virus genes, or uncharacterized.

## Data and code availability

All the scripts used for preprocessing and analysis are available in the GitHub repository https://lakhanp1.github.io/Pectobacterium_pangenome. Zenodo DOI for the GitHub repository: https://doi.org/10.5281/zenodo.13631140. Sequencing data can be accessed at NCBI with BioProjects [PR-JNA1122109](https://ncbi.nlm.nih.gov/bioproject/PRJNA1122109) for the NIVIP collection and PRJNA1140644 for the NAK collection.

## Supplementary information

**Additional file 1:** Supplementary tables.xlsx

## Ethics approval and consent to participate

Not applicable

## Consent for publication

Not applicable

## Competing interests

The authors declare that they have no competing interests.

## Funding

This research was funded by the Dutch Ministry of Economic Affairs in the Topsector Program “Horti-culture and Starting Materials” under the theme “Plant Health” (project number: LWV20.235) and its partners (NVWA, NAK, Naktuinbouw and BKD).

## Authors’ contributions

LP, SS, TL, DR: Conceptualization. LP: Data curation, Formal analysis, Investigation, Software, Writing – original draft. ID, MP: Resources. SS, TL, DR: Supervision, Funding acquisition, Project administration. LP, SS, TL, DR, AK: Writing - review and editing. All authors reviewed the manuscript. All authors have read and agreed to the published version of the manuscript.

## Acknowledgements

We would like to thank Harm Nijveen and the IT team of Wageningen University & Research for managing the Linux servers.

## Supplementary Figures, belonging to Pardeshi et al., “Pangenomics approach to understand the prophage dynamics in the radiating lineages of P. brasiliense”

**Supplementary Figure S1:**
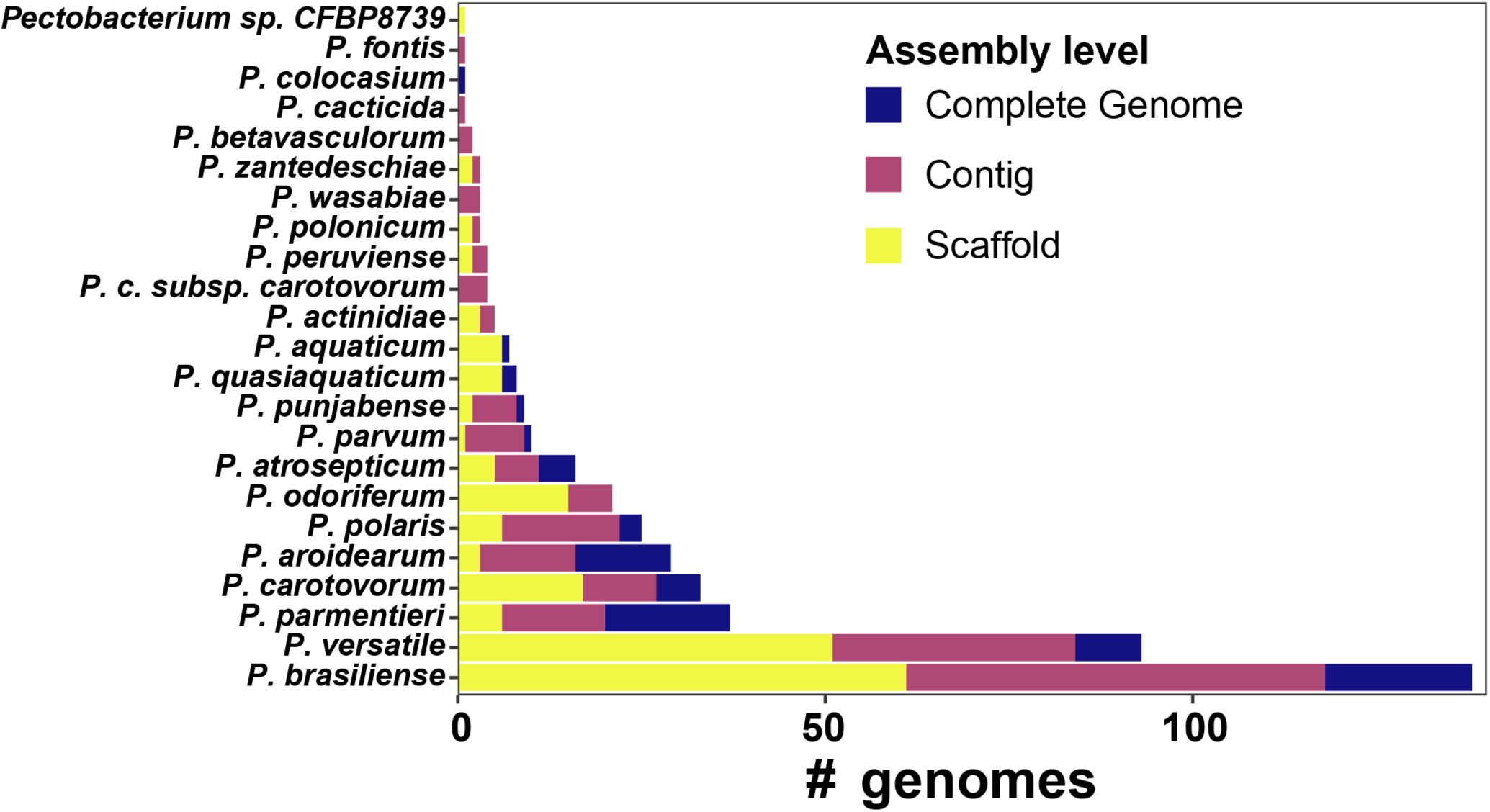
Numbers of *Pectobacterium* species genomes in the pangenome.

**Supplementary Figure S2:**
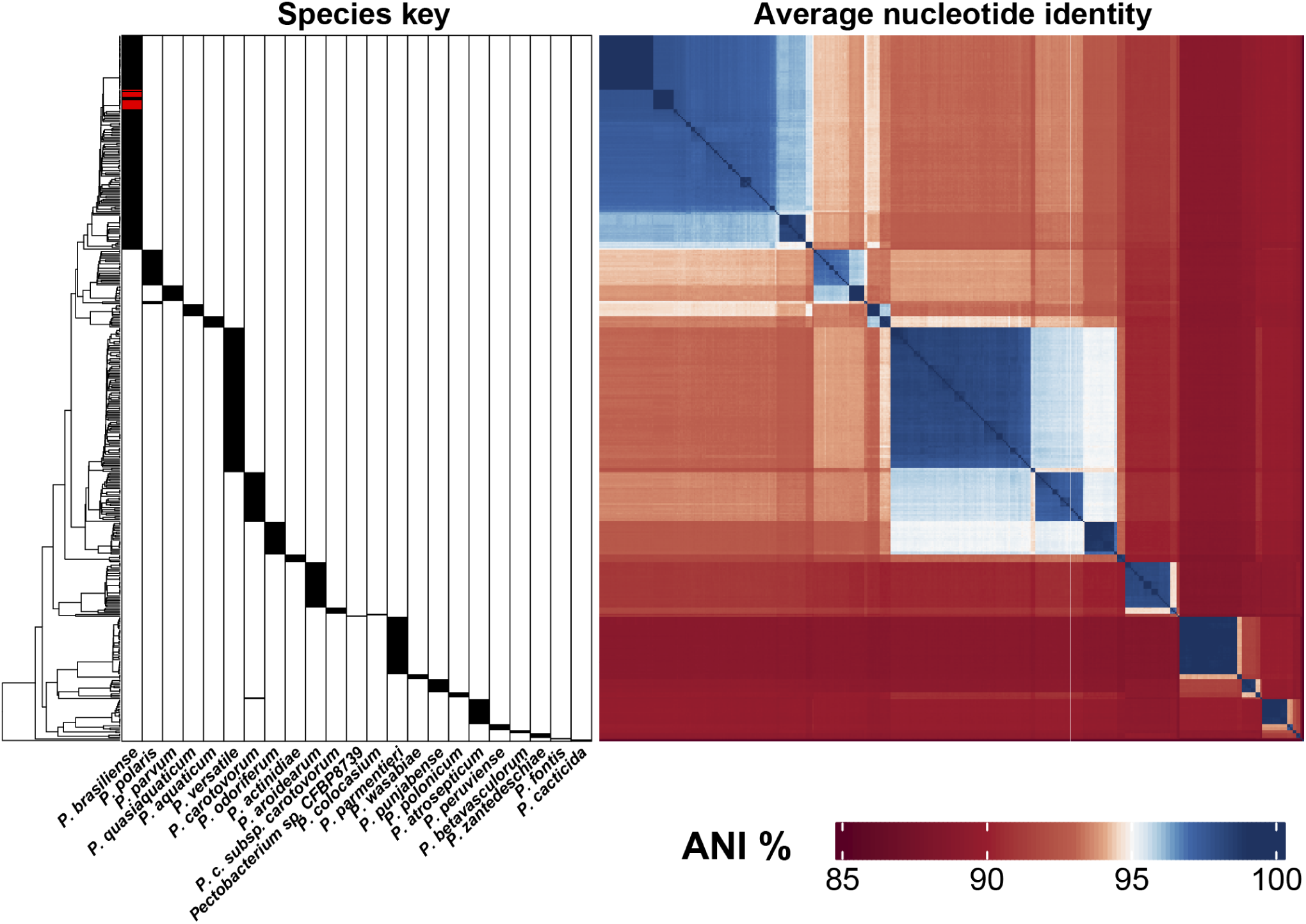
ANI between all pairs of *Pectobacterium* spp. genomes: from left to right, a dendrogram based on UPGMA clustering on a (1-ANI) distance matrix; species key, where FN-Pbr isolates are marked in red; ANI heatmap. ANI heatmap color scale is centered around white color which represents 95% ANI, a commonly used species delineation cutoff.

**Supplementary Figure S3:**
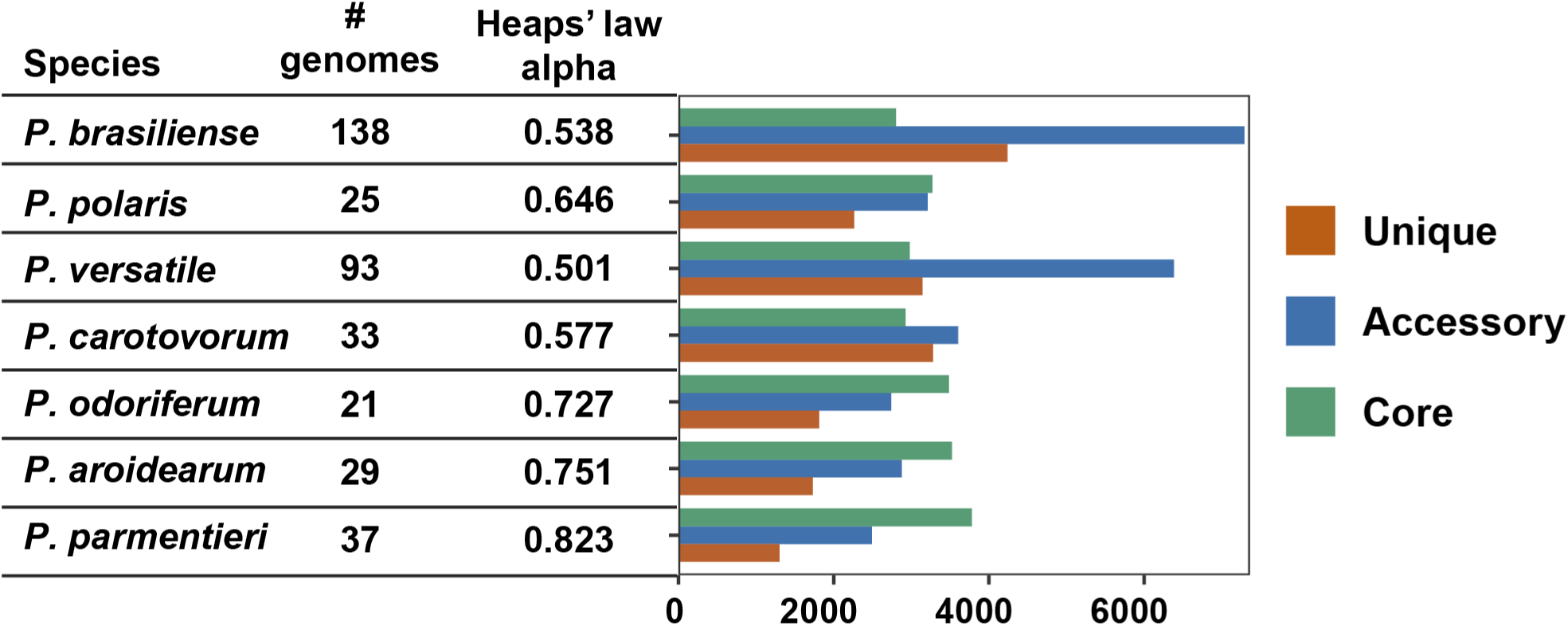
Core, accessory and unique gene statistics for *Pectobacterium* species, with pangenome openness depicted by Heaps’ law *α* value.

**Supplementary Figure S4:**
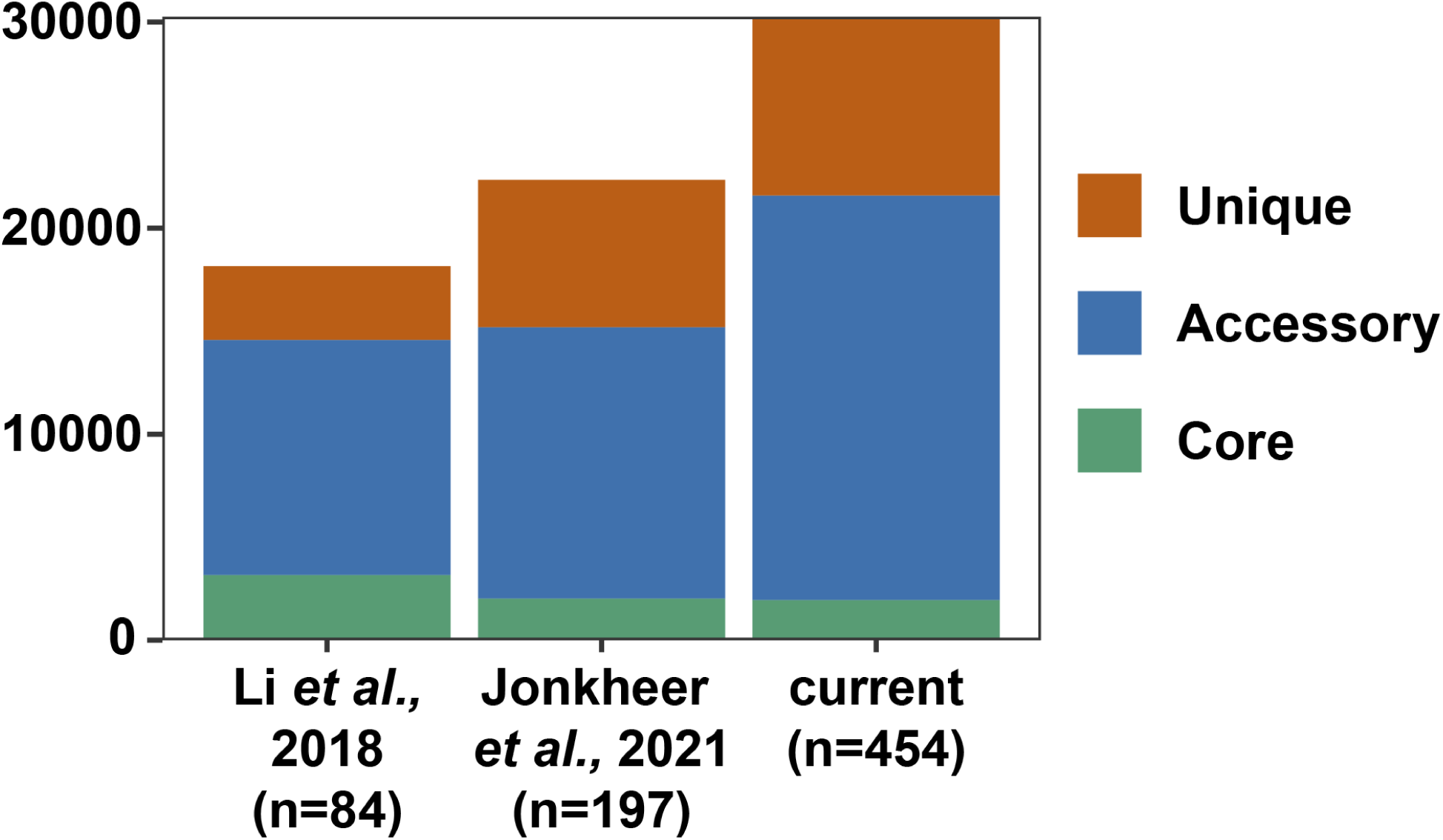
Homology group counts in previous *Pectobacterium* spp. pangenomes and the current one. Number of genomes in the pangenome are mentioned in the parenthesis.

**Supplementary Figure S5:**
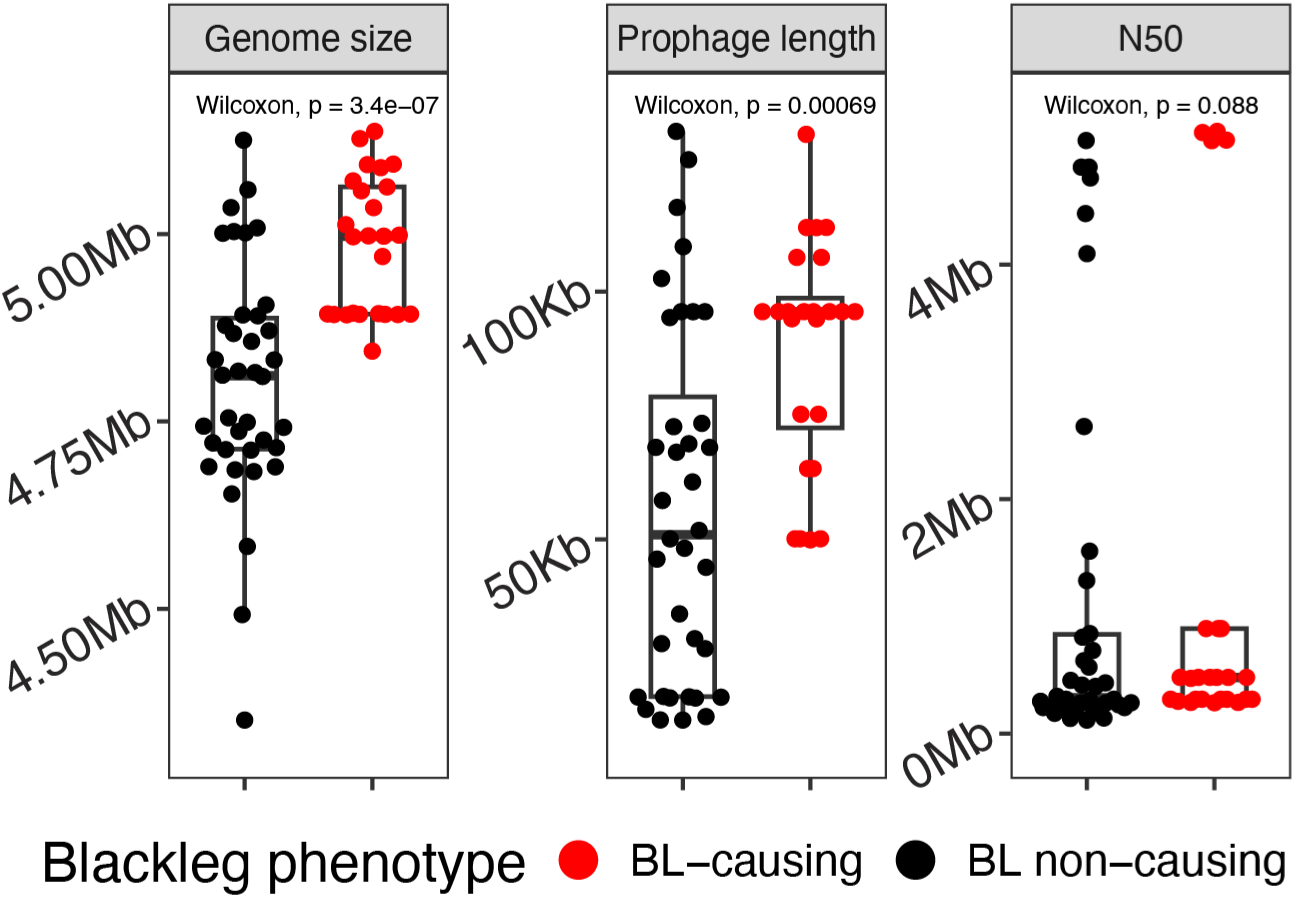
Prophage length comparison between BL-causing and non-causing *P. brasiliense* isolates

**Supplementary Figure S6:**
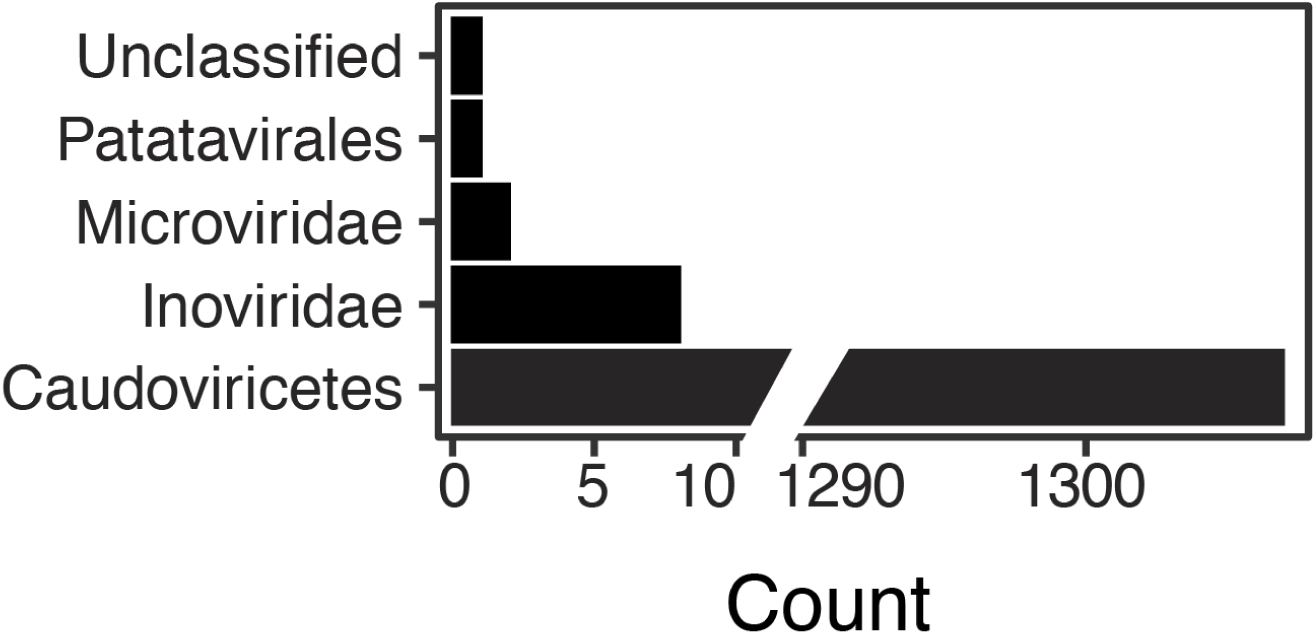
Prophage taxonomy

**Supplementary Figure S7:**
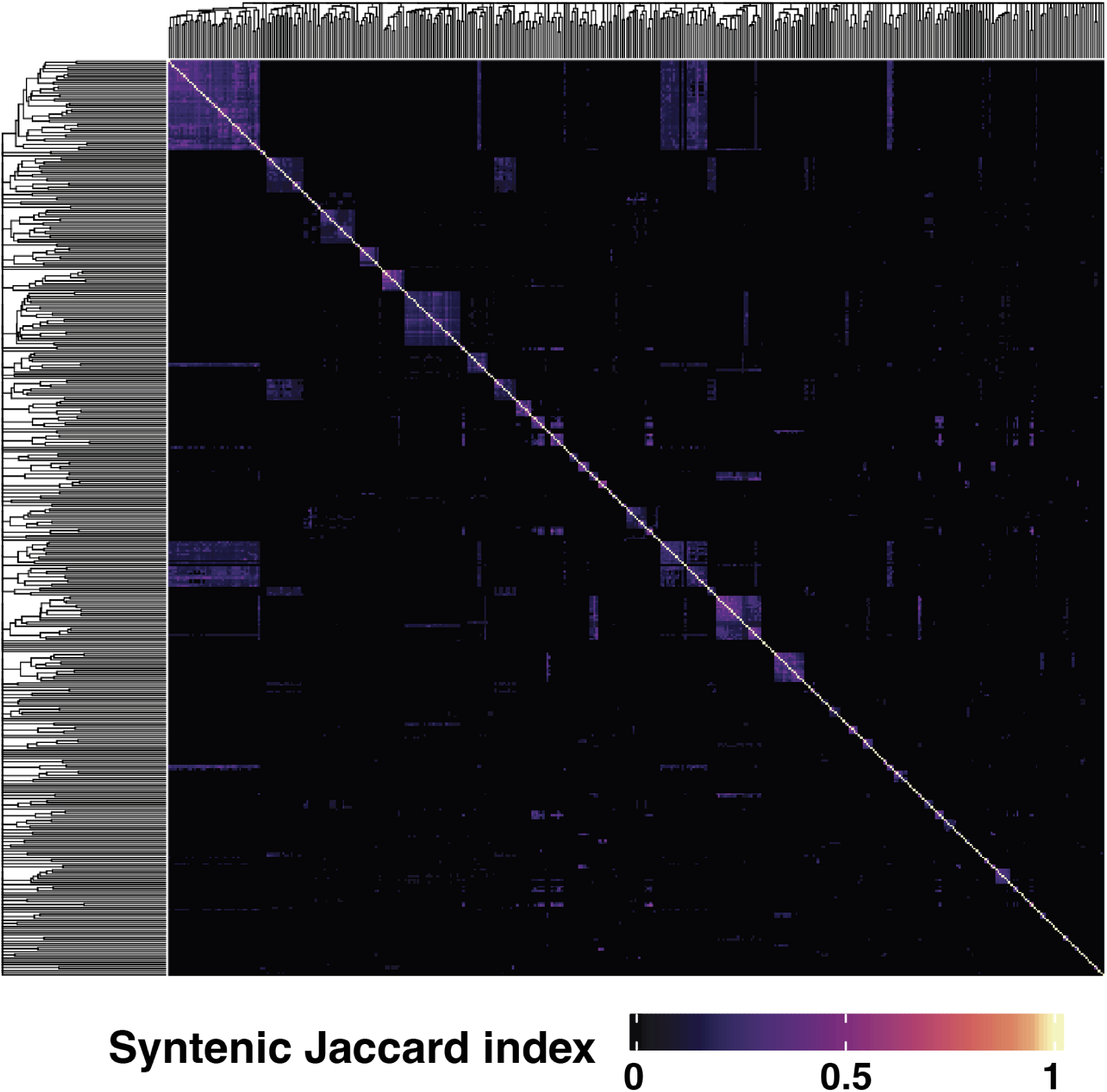
Syntenic Jaccard index heatmap for 436 representative prophages

## Notes

https://lakhanp1.github.io/Pectobacterium_pangenome

### Competing Interest Statement

The authors have declared no competing interest.

### Summary of Updates

Change the Zenodo DOI link; author order in PDF corrected.

